# gen3DNet: An R Package for Generating 3D Network Models

**DOI:** 10.1101/2024.11.11.623060

**Authors:** Paul Morrison, Tina Tang, Charles Lu, Shamim Mollah

## Abstract

**Motivation:** Networks are ubiquitous to organize and represent data in the real world such as the biomedical networks, social networks, etc. Due to the attractive property of networks which can explicitly capture the rich relationships among data samples, there is an increasing demand to transform linkage-free data to graph-structure data for efficient downstream analyses. Existing works typically focus on representing a single data object like gene expression profile in a homogeneous 2D network, which fail to deal with situations where two different data objects are involved to create a 3D heterogeneous network.

**Results:** In this paper, we introduce an R package, gen3DNet (a generic version of the iPhDNet), for generating 3D network models from two correlated objects with shared common factors. Specifically, gen3DNet builds the relationships between samples and shared factors using the non-negative matrix factorization, where three clustering techniques are evaluated to determine the number of functional modules. In addition, it builds the relationships between samples from the two distinct data objects based on a partial least squares regression model. Usage of the package is illustrated through a real-world application.

**GitHub URL** Source code is available at https://github.com/mollahlab/gen3DNet

## Introduction

Many real-world data and complex systems are in the form of networks, such as protein-protein interaction networks, gene regulatory networks, document networks, etc The structure and function of complex networks. The structure of scientific collaboration networks [1–4]. A typical network model can be represented as *G* = (*V, E*), where *V* ∈ *R*^*n*^ is a set of *n* unique vertices and *E* ∈ *R*^*n*×*n*^ is a set of edges capturing the pairwise relationships among vertices. Compared with the plain linkage-free data, data organized in networks have many attractive merits. First, the complex relevance and dependency relationships between data instances can be explicitly reflected by the edge connections. In addition to the explicit relations, networks are able to capture rich intermediate or implicit correlations between instances through the relationship propagation over the entire topology structure, *i*.*e*., two vertices with similar structure tend to be relevant. Another appealing factor is that data represented in networks are beneficial for efficient downstream applications, e.g., clustering, classification, prediction and visualization tasks, with the support of many off-the-shelf network-based learning algorithms [5, 6]. As a result, the past few years have seen a plethora of works which seek to transform linkage-free data to the equivalent network-structured data (a.k.a network construction). Specially, network construction has attracted tremendous attention in the biomedical domain. For examples, gene regulatory network construction aims at modeling the complex regulatory activities that may occur among genes by exploring the gene expression data [4, 7–9]. Gene co-expression network construction aims to build the pairwise transcript–transcript associations among genes by calculating their co-expression values [10]. Protein-protein interaction network construction aims to build the high-confident correlation relationships among proteins in various organisms using the high-throughput techniques [11, 12]. One common point for these network constructions lie in that they try to derive a 2D and homogeneous network model from the single data object such as gene expression and protein-protein interaction observations, which fails to consider situations in which two distinct correlated data objects are involved to create a 3D heterogeneous network. For instance, in the complex biological system of breast cancer microenvironment, protein regulations and histone regulations in the extracellular matrix can be viewed as two different data objects which are causally mediated by the shared growth promoting factors. Analogically, in the service computing domain, Mashups and APIs can be viewed as two different objects which are associated with the shared functional category factors. In general, the challenge of constructing a 3D network model from two different but correlated data objects is twofold: 1) the relationship characterization between the data object and shared factor; and 2) the relationship characterization between the two distinct data objects in a unified network model.

Here, we extends a recently proposed network model iPhDNet [4] (e.g., capturing the interactions among peptides, histons and drugs) to introduce **gen3DNet** encapsulated in an R package for generating 3D network models from the two input data objects with shared relevant factors. **Figure 1** shows the high-level implementation scheme of the proposed gen3DNet which takes two data objects as inputs. First, it performs the clustering on the first data object, e.g., using Non-negative Matrix Factorization (NMF) [13], to obtain the linkage coefficients between samples from the first object and shared factors, where *K* clusters (e.g., *C*_1_, *C*_2_, ⋯, *C*_*k*_) are generated for all samples and shared factors respectively. We implemented three methods to determine the optimal selection of *K*, including NbClust [14], Cophenetic [15], Silhouette [16] based techniques and evaluation metrics. Meanwhile, the linkage coefficients between samples from the two different objects are obtained based on the Partial Least Squares Regression (PLSR) model [17]. Finally, the two interaction networks generated in above two steps are integrated to a single 3D network.

**Figure 1.**
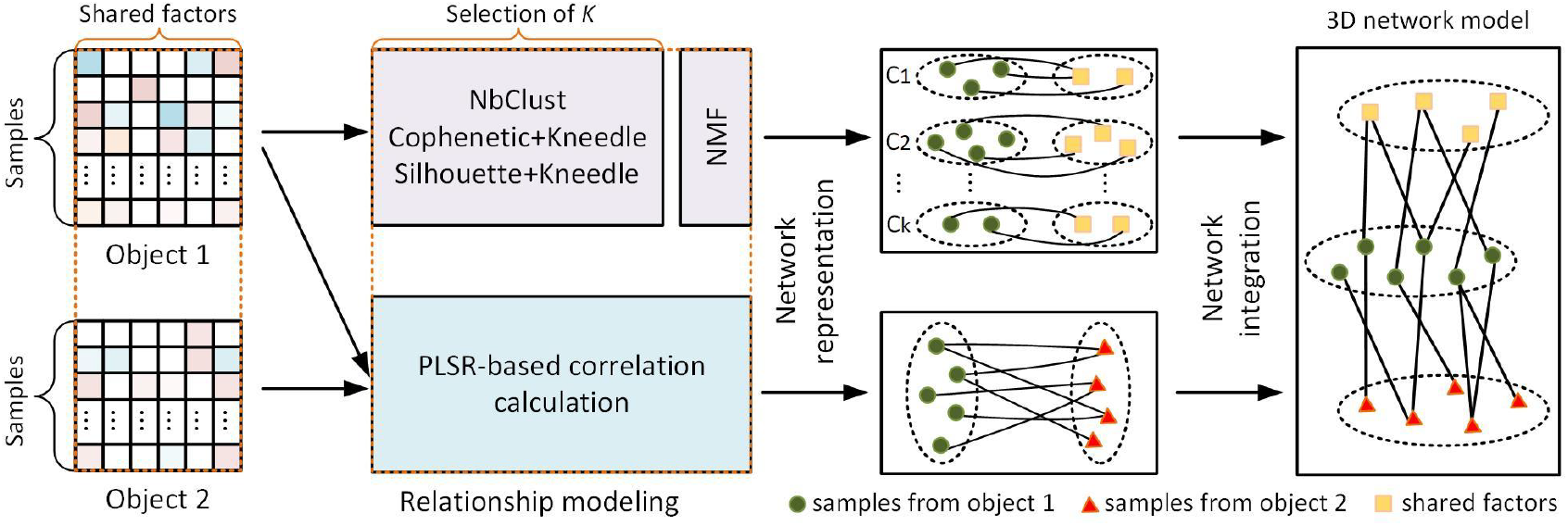
Implementation framework of the proposed gen3DNet.

The rest of paper is organized as follows. In the second section, we briefly describe the network modeling based on NMF and PLSR techniques. In the third section, we present the gen3DNet package with examples. In the forth section, we discuss the analysis results of a real-world application in the bioinformatics area based on gen3DNet. Finally, the last section concludes the paper.

## Materials and Methods

Let the two input data objects be represented as *M*_1_ ∈ *R*^*m*×*f*^ and *M*_2_ ∈ *R*^*n*×*f*^, where *m* and *n* are respectively the numbers of samples from the first and second objects, and *f* is the number of shared factors between the two objects. First, the relationships between the samples and shared factors are calculated from the first data object *M*_1_, which involves a non-negative matrix factorization is performed as:

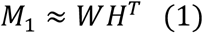

where samples and shared factors are mapped in a *k*-dimensional latent space represented in *W* ∈ *R*^*m*×*k*^ and *H* ∈ *R*^*n*×*k*^, respectively. *k* is a hyper-parameter which can be determined by the three evaluation methods including NbClust, Cophenetic and Silhouette shown in **Figure 1**. It is necessary to note that in order to guarantee the non-negativity of the input matrix in NMF, *M*_1_ has been previously log-transformed which is a prevalent practice in the biomedical domain. Then, all samples are grouped into *k* clusters based on *W* in which each sample is assigned with a cluster with the largest membership value across the *k* latent dimensions, *i*.*e*., each of the *k* latent space dimensions corresponds to a cluster. The same clustering process applies to all shared factors based on *H*. Finally, samples and shared factors from the identical clusters build the pairwise edge connections, where the weights on the edges are given by the corresponding coefficient values in *H, i*.*e*., assume sample *s*_1_ and factor *p*_*j*_ are in the *k*^*th*^ cluster, then the connection weight between *s*_1_ and *p*_*j*_ equals to *H*_*j,k*_.

Then, the relationships between samples from the two different objects are calculated based on the PLSR model:

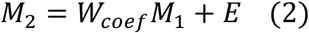

where *W*_*coef*_ ∈ *R*^*n*×*n*^ is the coefficient matrix revealing the contributions of the first object *M*_1_ to the formulation of the second object *M*_2_, and *E* is the residual or error matrix for the PLSR model. We then build the connection between samples from object 1 and samples from objects 2, where the edge weights are given by *W*_*coef*_. By integrating the output networks from the NMF and PLSR modelings, a 3D network model can be finally produced shown in **Figure 1**.

### Default k selection algorithm

An important component of iPhDNet [4] is the number of clusters present within the data, but there is no single domain-based method for choosing this parameter. For this reason, we undertook an evaluation of some common k-selection algorithms to determine which can be expected to generalize best. To do this, we began by generating hundreds of synthetic datasets, each created to have a specific number of clusters, ran the candidate algorithms on these datasets, and observed the correlation between the true and predicted k values under varying conditions. Based on these results, we chose a combination of the Ward clustering algorithm and KL index (implemented in the NbClust library) as the default algorithm for k selection in gen3DNet.

Each dataset was generated by starting with k cluster centers, and then adding other data points in a Gaussian distribution around each center, with the standard deviation of this distribution being an important parameter. Another parameter was the number of dimensions (columns) used in the dataset. Both of these were varied using the following values:

- Options for stdev. within / stdev. between: 1, 3, 5, 7, 9, 11.
- Options for the true number of clusters: 2, 3, 4, 5, 6, 7, 8, 9, 10.
- Options for the number of dimensions: 2, 4, 8, 16, 32.

The number of samples (rows) in each artificial dataset was left constant (at 1000). Examples of sample datasets are shown in **Figure 2a, 2b**. Both have the same number of clusters, but this harder to tell in **Figure 2b** because the variation within clusters is similar to the variation between clusters.

**Figure 2.**
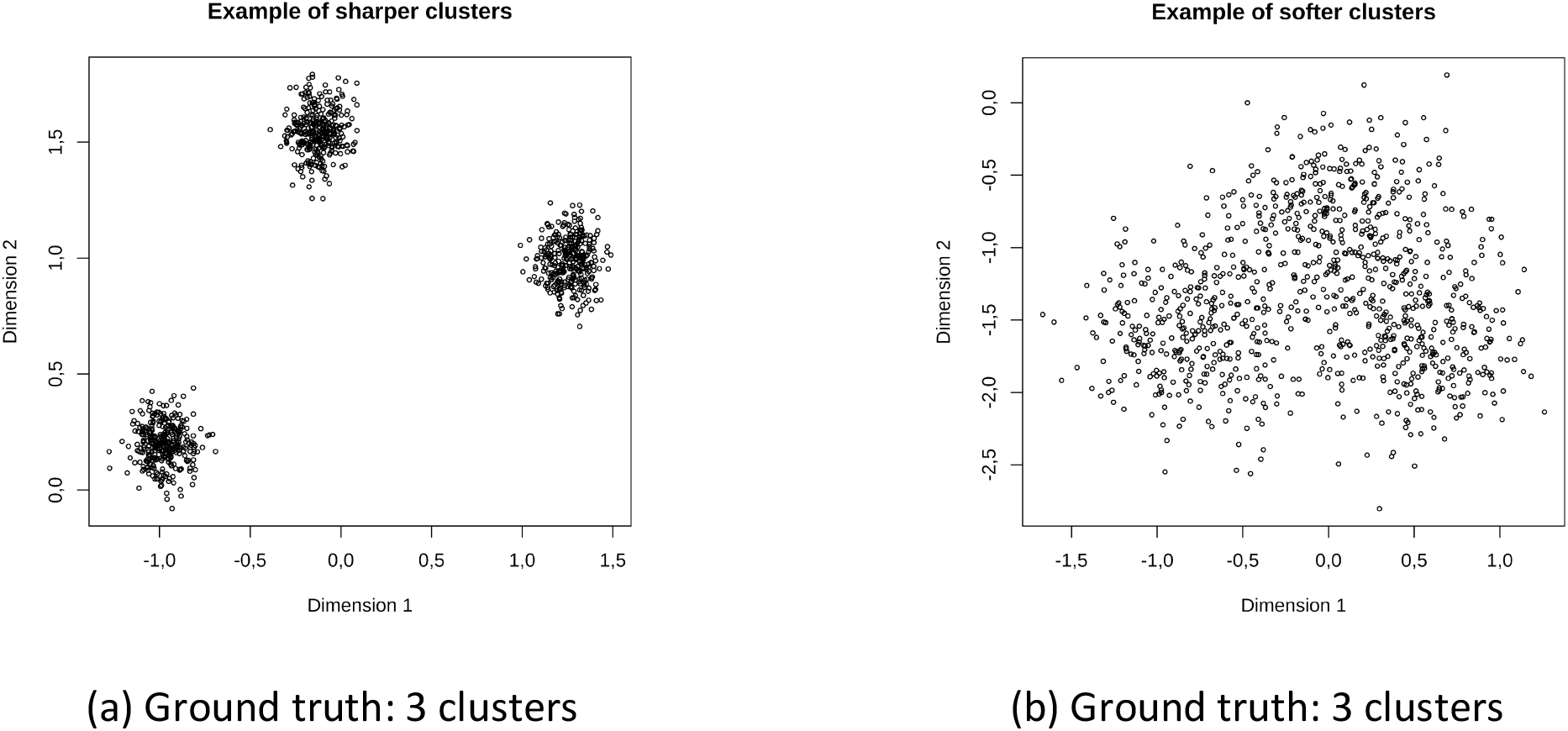
Example of simulated ground truth data.

The detailed dependency relations between modules in gen3DNet is depicted in **Figure 3**.

**Figure 3.**
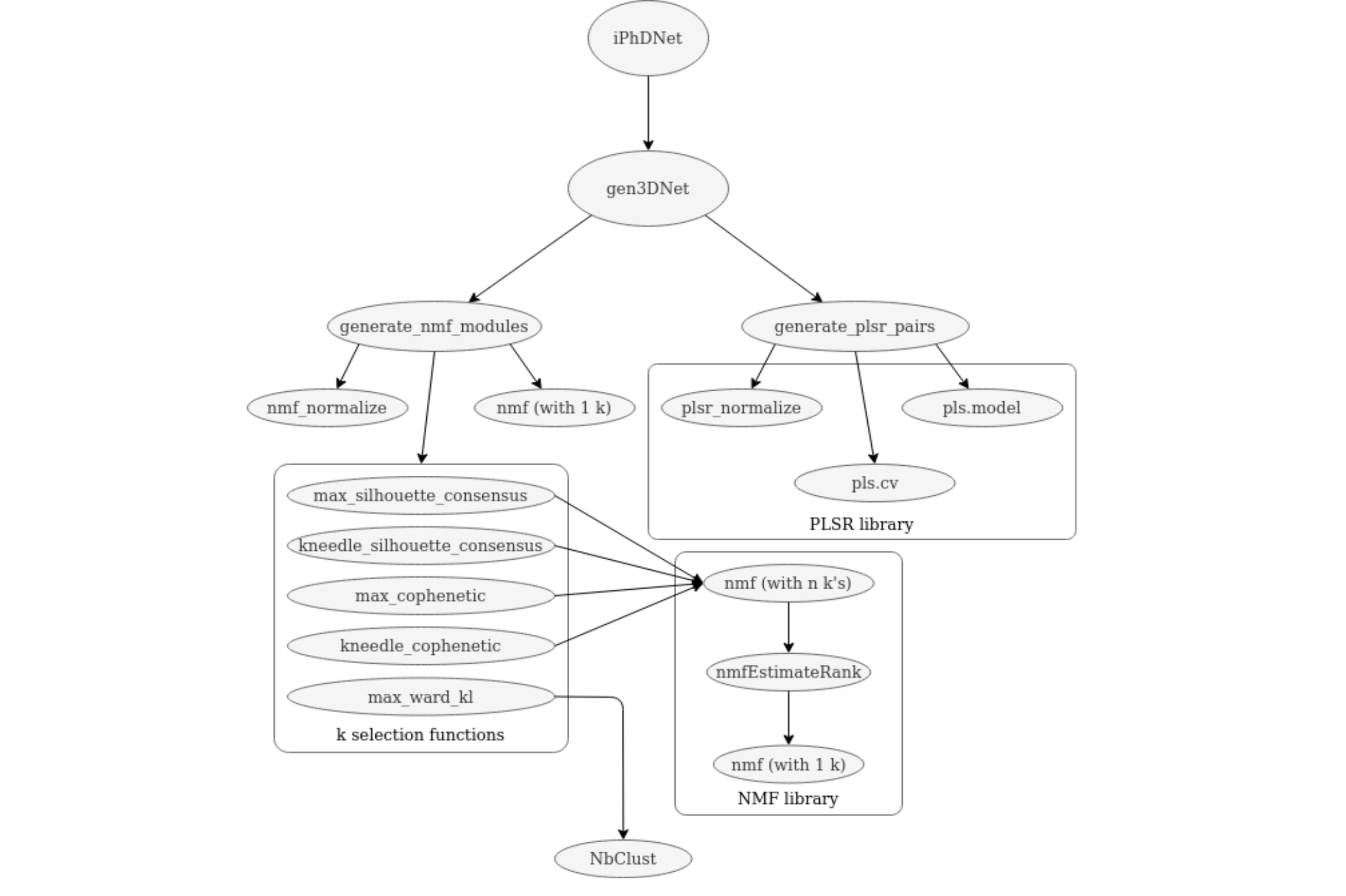
The dependency relations between modules in the gen3DNet package.

The following candidate k-selection methods were used:

#### 1. Cophenetic correlation

Given a hierarchical classification, the cophenetic correlation is the correlation between the true distances and dendrogrammatic distances of each pair of data points.

As the number of clusters increases, the cophenetic correlation continues to improve, so choosing the maximum will favor the maximum number of clusters. For that reason, a separate algorithm is needed to find the “knee", where increasing k any further has no benefit. For this purpose, we used the Kneedle algorithm [18]. Since the knee often appears in an “L” shape, the Kneedle algorithm works by drawing a line from the upper left to the lower right and locating the point that is furthest from this line. This leads the algorithm to the bend of the “L".

#### 2. Silhouette

defines the silhouette for an individual point as a number from -1 to +1 comparing the closeness of other points within the same cluster to the rest of the points in the dataset. For a point i, let a(i) be the average distance between i and the other points in its cluster, and b(i) be the average distance between i and the closest cluster to it. The silhouette for this point is

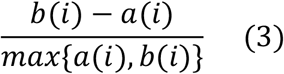

For an entire dataset, the silhouette is the average over all points. This method was also combined with the Kneedle algorithm for knee detection.

#### 3. Ward KL

This k-selection method uses the Krzanowski-Lai index [19] to choose the number of clusters, with the cluster assignments being made by Ward hierarchical clustering algorithm [20].

The Ward algorithm, described in, is a hierarchical clustering algorithm that minimizes a measure of within-cluster variance (in this case, the squared Euclidean distance between points). Initially, each point begins as a separate cluster. In each iteration, a pair of clusters must be merged - and the pair with the lowest within-cluster distance is always the one chosen. The “ward.D2” version of the Ward algorithm, from the NbClust library, was the implementation used.

To turn this clustering method into a k-selection algorithm, the NbClust library finds clusters for each number from 1 to 32 and picks the one with the highest score. In this case, the score used for the decision was the index proposed in.

The Krzanowski-Lai index is based on the quantity 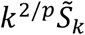, where p is the number of columns and 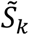 is the trace of the pooled within-group covariance matrix. The difference between a given value of k and the previous value is then referred to as *DIFF*_*k*_.

Finally, the Krzanowski-Lai index itself is defined as

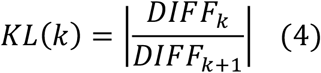

Therefore, the KL index will be maximized when the change from adding another cluster (|*DIFF*_*k*+1_|) is the smallest compared with the change from adding the last cluster (|*DIFF*_*k*_|).

## The gen3DNet package

We encapsulate above modeling process in the R package gen3DNet which is available from the CRAN at https://CRAN.R-project.org/package=gen3DNet.

### Installation

The package can be installed in two ways.

1) The release version from CRAN:

install.packages(“gen3DNet”)

or 2) Directly from GitHub:

install.packages(“gen3DNet”) remotes::install_github(“MollahLab/gen3DNet”)

## Experiments and Results Analysis

### Test cases

In this section, we demonstrate an application of gen3DNet to build a 3D network model linking phosphoproteins, histones and drugs data obtained from Mollah SA and Subramaniam S study [4] and compared the results with iPhDNet. Gen3DNet results are in perfect agreement with iPhDNet results.

### Evaluation

To evaluate Gen3DNet, 20 datasets were generated/simulated for each combination of (n_rows, stdev. within / stdev. between). Each of the three k-selection methods was applied to the datasets. The results for one pair of settings are shown in **Figure 4**, which shows the true k on the x axis and the predicted k on the y axis. As the figure shows, the correlation between the true number of clusters and the Ward prediction is perfect (45 degree angle line), while it is weaker for the other methods: Kneedle Silhouette and Kneedle Cophenetic (scattered throughout).

**Figure 4.**
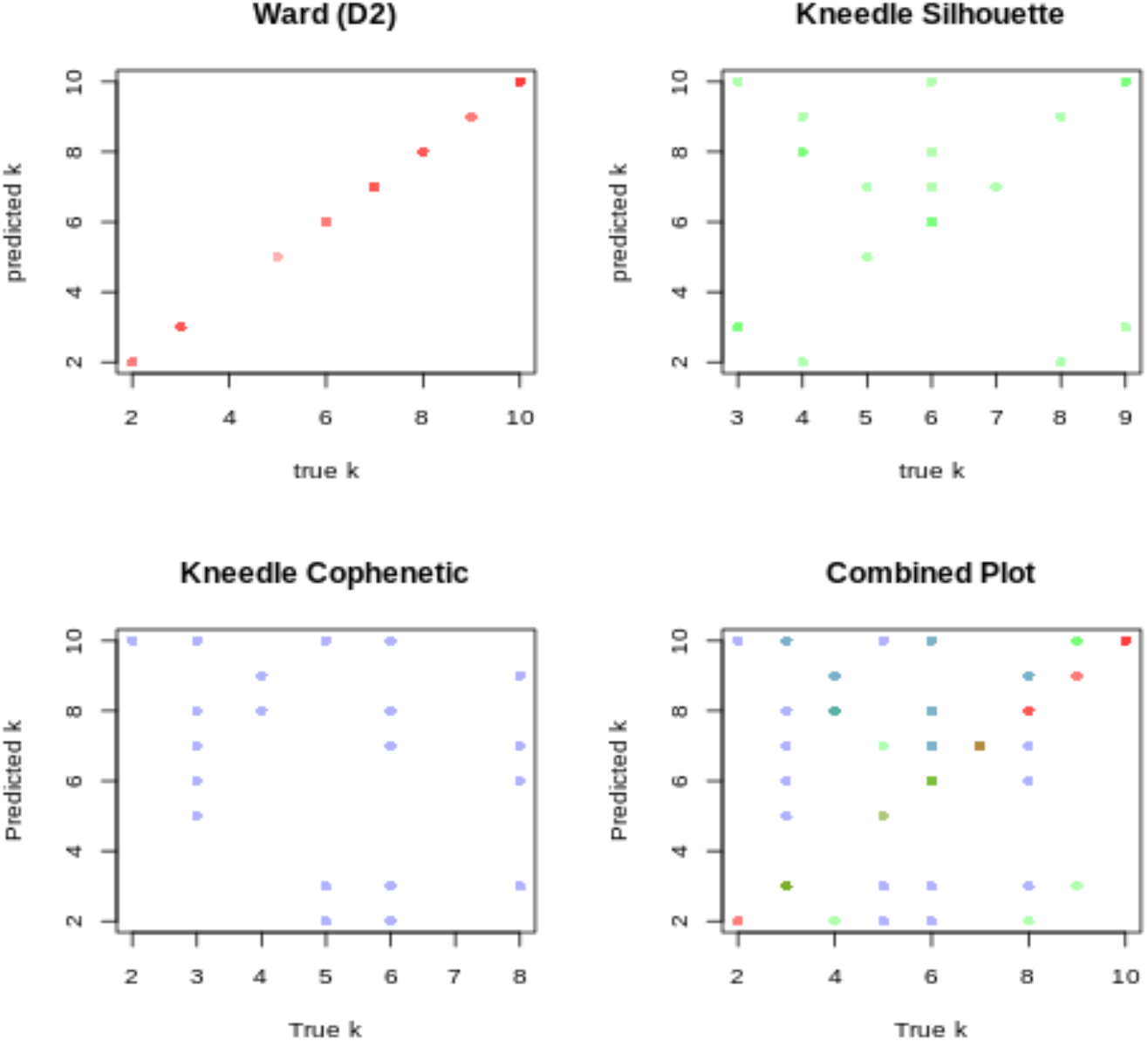
Comparison of the three adopted methods for identifying the optimal number (K) of clusters.

To compare the performance of these clustering methods across different settings, each of these plots was represented as a correlation - and the results were gathered into heatmaps in **Figure 5**. From these figures, we see that the Ward method improves as the number of dimensions increases and the clusters become sharper. This makes sense, as either change will increase the distance between clusters. This pattern is not seen for the other k-selection methods.

**Figure 5.**
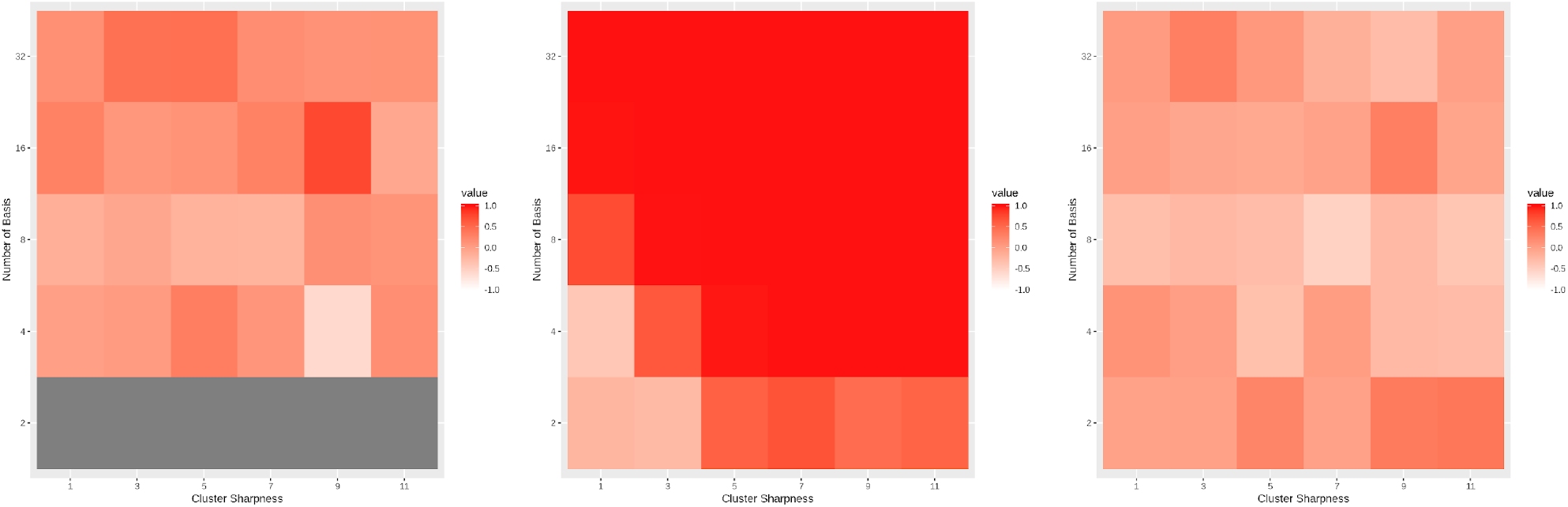
Average correlation in random datasets with different cluster sharpness and basis.

## Discussion

The past few years have seen an increasing demand to transform and represent linkage-free data in networks, which will deliver higher efficiency for many downstream applications. In this paper, we introduced a simple and generic tool for generating 3D network models from two input matrices with shared column properties. We have demonstrated the usage of the proposed gen3DNet R package and its application to construct a 3D network involving phosphoproteins, histones and drugs in the breast cancer study. Beside studying the histone signatures and precise mechanism of action in cellular reprogramming of histone modifications in breast cancer, gen3DNet can be widely used to reveal 3D connectivity and shared associations within any complex systems.

## Acknowledgments

This work was supported by Washington University in St. Louis School of Medicine Genetics and Institute for Informatics and Data Sciences departmental funds.

## Author Contributions

Original concept: S.M.; experimental design: SM; implementation: P.M., S.M.; Evaluation: P.M., S.M.; manuscript writing: P.M., S.M. R package submission: T.T., C.L.

